# The Decay of Disease Association with Declining Linkage Disequilibrium: A Fine Mapping Theorem

**DOI:** 10.1101/052381

**Authors:** Mehdi Maadooliat, Naveen K. Bansal, Jiblal Upadhya, Manzur R. Farazi, Zhan Ye, Xiang Li, Steven J. Schrodi

## Abstract

Several important and fundamental aspects of disease genetics models have yet to be described. One such property is the relationship of disease association statistics at a marker site closely linked to a disease causing site. A complete description of this two-locus system is of particular importance to experimental efforts to fine map association signals for complex diseases. Here, we present a simple relationship between disease association statistics and the decline of linkage disequilibrium from a causal site. A complete derivation of this relationship from a general disease model is shown for very large sample sizes. Quite interestingly, this relationship holds across all modes of inheritance. Extensive Monte Carlo simulations using a disease genetics model applied to chromosomes subjected to a standard model of recombination are employed to better understand the variation around this fine mapping theorem due to sampling effects. We also use this relationship to provide a framework for estimating properties of a non-interrogated causal site using data at closely linked markers. We anticipate that understanding the patterns of disease association decay with declining linkage disequilibrium from a causal site will enable more powerful fine mapping methods.

## Introduction

Genetic markers closely linked to disease-causing sites will exhibit association with disease through linkage disequilibrium.^1-4^ This is the central idea behind population-based association mapping of disease genes using high density SNP arrays.^5,6^ However, the decay of disease association with declining linkage disequilibrium from a disease-predisposing, functional site has not yet been completely described even though this is a fundamental property of disease genetics. Doing so will provide much needed information concerning the properties of disease genetics and greatly aid experimental designs and statistical methods for identifying functional variants in regions that exhibit disease association.

Although many have argued that genome-wide association studies have been largely unsuccessful in that they have not revealed a large proportion of the heritability from most complex diseases,^7^ it is certainly clear that numerous loci with impressive statistical evidence for correlation with a wide variety of complex diseases have been identified and replicated.^8^ In a number of instances, these results have provided much needed insight into the biochemical pathways and cellular mechanisms responsible for increasing disease risk.^9-12^ However, the functional variants underlying the majority of these disease-associated regions have yet to be identified and fully described.^13^ The dearth of information concerning functional variants obviously presents a sizable impediment to further dissection of complex disease etiologies. If genetic and statistical methods can aid in generating either supporting or opposing evidence for the role of functional motifs within a region of association, then the progression of human genetics studies can be made much more efficient and potent.

When designing fine mapping genotyping experiments, it is important to select genetic variants and subregions so that adequately cover two types of disease models are adequately covered (i.e. the fine mapping design is well-powered to discover the functional variants). The first class of model that should be covered by such efforts would be models of a causal variant driving a portion, or perhaps all of the disease association within a region. Under this model, varying levels of association signal at different sites are explained by different levels of linkage disequilibrium with causal variants. Hence, given allele frequencies and linkage disequilibrium patterns, one can, in principle, back-calculate the properties of putative functional variants that could be driving an initially observed disease association within the region of interest. Known variants, including those that were not initially interrogated, fulfilling these calculated allele frequency and linkage disequilibrium properties with the initial markers should then be included in a fine-mapping panel. The second model to be covered by a fine-mapping panel of markers is one of allelic heterogeneity at a functional motif (e.g., a gene) that was originally found to exhibit a disease association signal. Empirical data tends to strongly favor this type of model over an individual variant serving as the sole driving allele within a region.^14-18^ Indeed, it is quite typical for studies aiming to fine map regions harboring a GWAS-significant SNP to reveal multiple disease-correlated variants within the same gene. This is not terribly surprising as the site frequency spectrum is expected to contain vast numbers of rare variants in outbred populations, which is accentuated in rapidly expanding demographics.^19-21^ Even if there is a small likelihood of any one of these rare variants to exhibit pathogenic effects, the sheer number of variants segregating at a gene trends to produce multiple functional alleles. To cover this class of disease models, one would want to reliably identify the functional motifs tagged by an initial association signal and proceed by exhaustively interrogating variants within those functional motifs. In practice, this two-model approach guiding fine mapping was employed successfully to identify alleles segregating at the *TRAF1-C5* region conferring susceptibility to rheumatoid arthritis.^22,23^

To address the statistical aspects of prioritizing potentially causal variants within a fine-mapped region, several methods have been developed including a useful Bayesian method was developed by Maller and colleagues,^24^ which uses Bayes Factor for each variant in the region and calculates the proportion of the total sum of Bayes Factors in the region that is attributable to that variant, producing a relative ranking of the strength of evidence for each variant within the disease-associated region being causal. These calculations allow for the determination of a credible set of highest ranked variants that explains the large majority of the statistical evidence of disease association within the region of interest. The Maller et al. method has been applied to fine mapping data for complex diseases, such as type 1 diabetes.^25^ Other important developments in fine mapping approaches include another Bayesian approach, Bim-Bam^26^, methods which determine subsets of variants that likely contain causal sites, CAVIAR^27^ and CAVIARBF^28^, coalescent-based methods^29-31^ and PAINTOR^32^, which incorporates functional annotation data in a probabilistic manner.

Here, building upon previous work,^3^,^33-36^ we prove a simple, analytic relationship between case/control association statistics at two closely-linked sites and the linkage disequilibrium between the two sites under a generalized disease genetics model. The result holds for very large sample sizes. Interestingly, the result is invariant with mode of inheritance parameters. Further, we posit that concurrently considering the patterns of disease-association and the genetic architecture within a region of interest may strengthen the ability to assess the likelihood that a particular variant is indeed causal with regard to inflating the risk of disease. By doing so, one may be better able to prioritize variants for functional follow-up studies. For finite sample sizes, dispersion around this relationship is expected and we therefore explore this variation in the result through the use of a Monte Carlo simulation.

## Approximation

Under the Pritchard-Przeworski derivation^33^, the power to detect disease association at a causal site and marker site were found to be approximately the same if the sample size at a marker site is increased by a factor of (*r*^2^)^−1^ over that used in interrogating the causal site. *r*^2^ is the standard measure of linkage disequilibrium between the causal site and the marker site. While certainly an intriguing relationship between sample sizes, as it is, the finding may not always have utility in fine mapping applications as most association studies use the same number samples at all sites interrogated. That said, this relationship can be used to motivate related and illuminating properties regarding how fast the disease association signal can be expected to decay as a function of declining linkage disequilibrium from a causal site. Equating the power at the disease-predisposing site to that at the marker site, it follows that

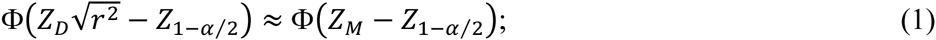

where *Z*_*D*_ and *Z*_*M*_ are the normally-distributed Z-scores for testing disease-association at the causal site and marker site, respectively; and α is the significance level. Taking the inverse functions and squaring yields the provocative approximation

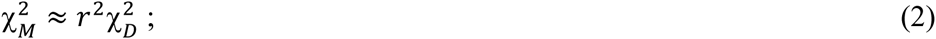

where 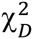 and 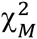 are the chi-squared statistics for disease association at the disease and marker sites, respectively. Plotting this approximation with the χ^2^ disease-association statistic on the ordinate and 1 – *r*^2^ on the abscissa is a simple method of displaying the expected linear decay in the χ^2^ values as the linkage disequilibrium with a causal site declines at different marker sites. **Figure 1** shows this relationship. This decay pattern was first used empirically in 2008 to fine map the *IL23R* region in psoriasis^37^ and has subsequently been used in other applications.^38^ Although this approximation is very useful in understanding the decay of disease association with declining linkage disequilibrium from a causal site, several simplifying assumptions were made in the original Pritchard-Przeworski derivation and therefore it is not known how violations of the original assumptions might produce departures from **Eqn 2**. Hence, an exact relationship between disease association statistics and *r*^2^ values with a causal site would aid in clarifying this relationship and motivate statistical approaches to harnessing this pattern for the purpose of fine-mapping functional alleles. Additionally, the allele frequencies are treated as parameters instead of random variables with sampling variances. So, understanding the dispersion around the decay patterns for finite sample sizes would further elucidate the relationships studied.

**Figure 1.**
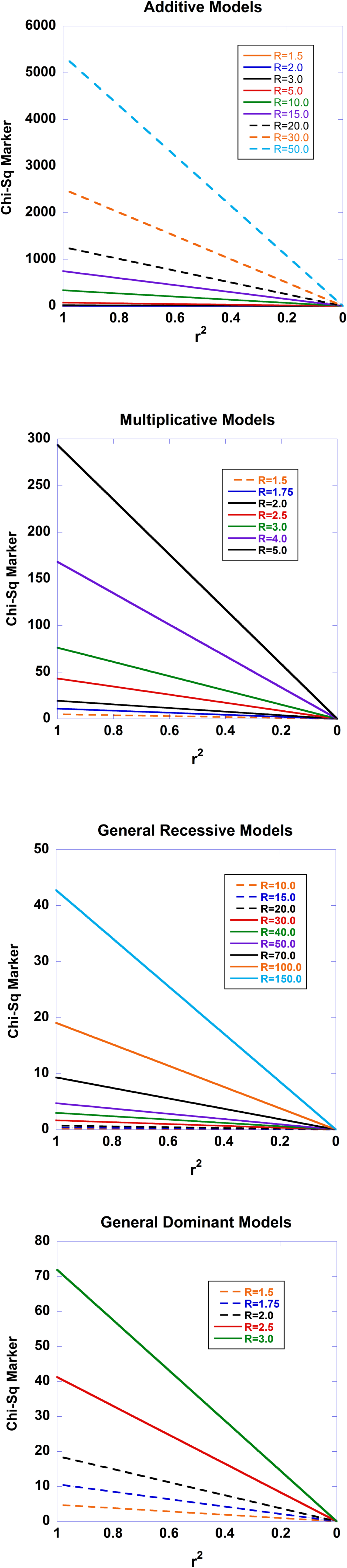
The Expected Decay of Disease Association with Declining Linkage Disequilibrium for Four Modes of Inheritance. The standard recursive haplotype frequencies under recombination were used to generate a series of haplotype combinations. The disease-predisposing allele at the causal site was set at a general population frequency of 0.01. The penetrance f_22_ was set to 0.001 and the remaining two penetrances varied according to the modes of inheritance examined and the relative risks cited in the Figures. Sample sizes were set at n_D_=2000 and n_C_=2000. **Fig. 1a** displays the results for an additive model, such that f_12_ is the arithmetic mean of f_22_ and f_11_. **Fig. 1b** shows the results under a multiplicative model. **Fig 1c** shows the results under a general recessive model. **Fig 1d** shows the results under a general dominant model.

## Full Derivation

Defining the Chi-Squared test statistics following the Pritchard-Przeworski derivation,

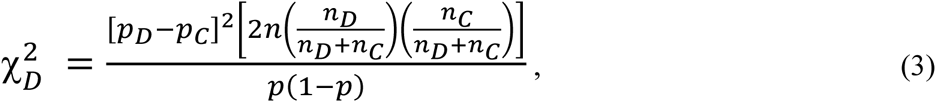

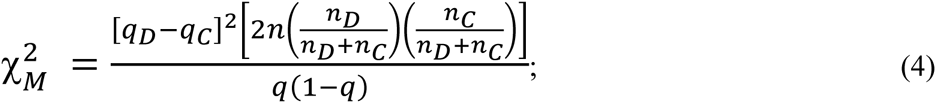

Where *p*, *p*_*D*_, and *p*_*C*_ are the frequencies of the *A*_1_ allele in the combined population, disease-affected population, and the control population, respectively and where *q*, *q*_*D*_, and *q*_*C*_ are the frequencies of the *B*_1_ allele in the combined population, disease-affected population, and the control population, respectively. *n*_*D*_ and *n*_*C*_ are the sample sizes for diploid cases and controls, respectively, and *n* = *n*_*D*_ + *n*_*C*_. For this work, we will assume that cases and controls are drawn from the general population such that the cases and controls are drawn with probabilities corresponding to the disease and healthy proportions.

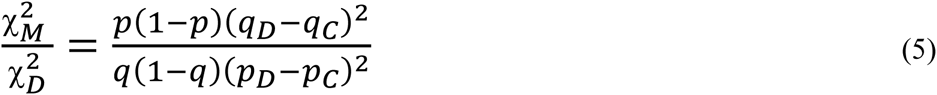

Noting that

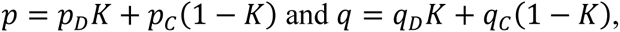

we can substitute 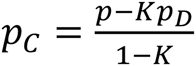 and 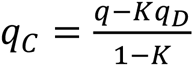 into **Eqn (5)**, resulting in

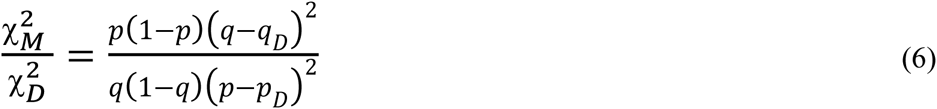

The next aim in the derivation is to substitute quantities for the allele frequencies in the affected population at both sites in terms of penetrances, disease prevalence, and general population allele frequencies,

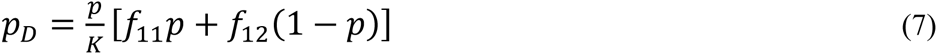

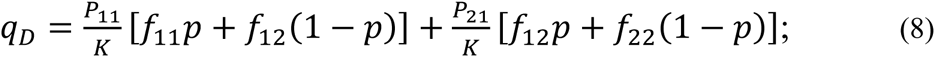

where *ƒ*_11_, *ƒ*_12_, and *ƒ*_22_ are the prevalences of the *A*_1_*A*_1_, *A*_1_*A*_2_, and *A*_2_*A*_2_ genotypes, respectively, such that *ƒ*_*ij*_ = *P*(*Case*|*A*_*i*_*A*_*j*_); *K* = *P*(*Case*), which, under this monogenic model and assuming Hardy-Weinberg Equilibrium in the general population, *K* = *ƒ*_11_*p*^2^ + 2*ƒ*_12_*p*(1-*p*) + *ƒ*_22_(1-*p*)^2^; and haplotype frequencies *P*_11_ = *P*(*A*_1_*B*_1_), and *P*_21_ =*P*(*A*_2_*B*_1_). Applied to complex diseases, it may be useful to think of this disease model as the subset of individuals with a common disease that is primarily driven by a particular locus. With the substitution into **Eqn 6**,

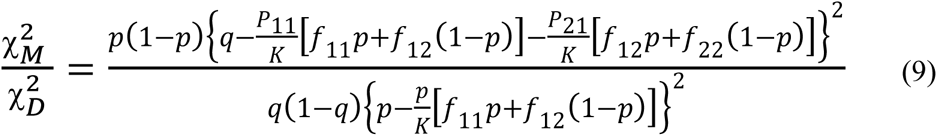

In **Eqn 9**, the R.H.S. numerator can be simplified to

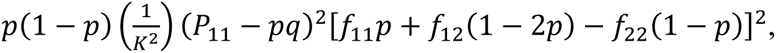

whereas the denominator in **Eqn 9** can be simplified to

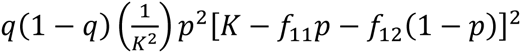

Hence, **Eqn 9** can be written as

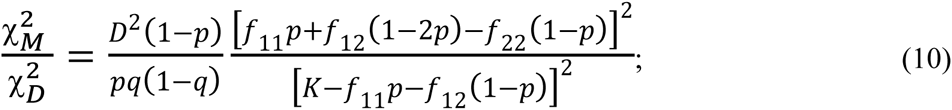

where *D* = *P*_11_*P*_22_ - *P*_12_*P*_21_ = *P*_11_ - *pq*.

Substituting *K* = *ƒ*_11_*p*^2^ + 2*ƒ*_12_*p*(1 - *p*) + *ƒ*_22_(1 - *p*)^2^,

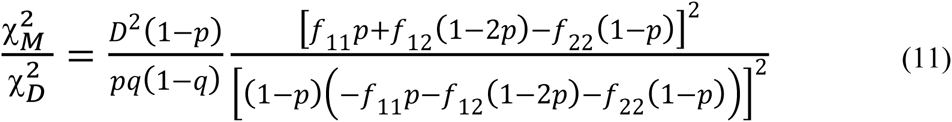

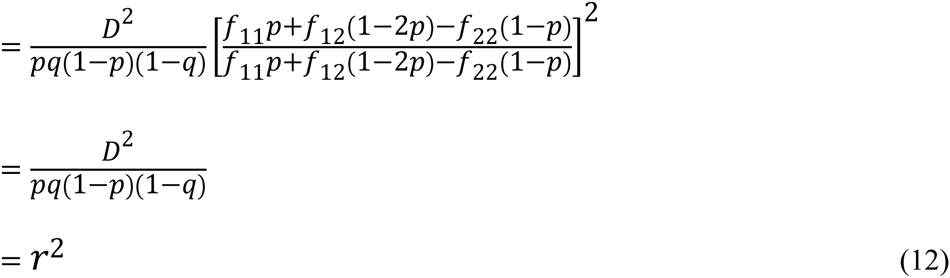

Therefore, we have shown the exact relationship under our model

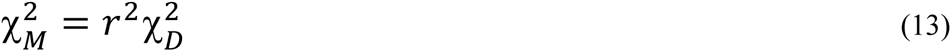

Not only is this relationship an exact result under the model employed, but it is universal in that there is no dependence on the penetrances. Thus, we may expect that from a true disease-susceptibility site, that there should be a linear decay in the Chi-squared statistics for disease association with declining *r*^2^ values with the causal site. **Figure 1** shows the expected disease association decay with declining linkage disequilibrium from the causal site for additive, multiplicative, recessive and dominant sets of models. The patterns arising from various relative risks are presented. Similarly, **Figure 2** presents the patterns expected as a function of sample sizes. Aside from **Eqn 13** illuminating a central aspect of disease genetics, we suspect that it carries utility in fine mapping applications – we hypothesize that identifying this type of pattern in fine mapping data will better enable the pinpointing of truly causal sites through harnessing correlated data.

**Figure 2.**
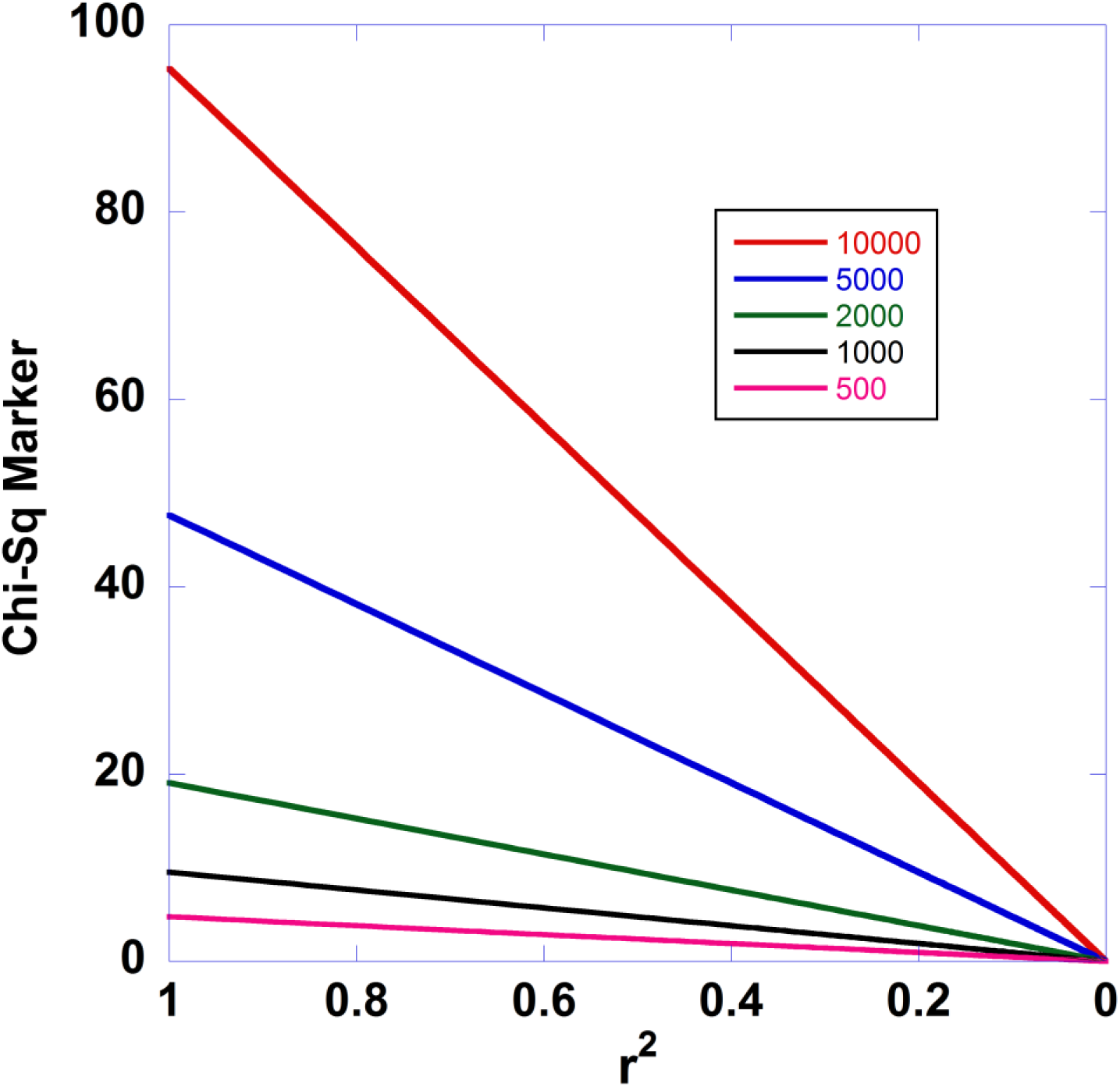
**Effect of Sample Size on the Expected Decay of Disease Association with Declining Linkage Disequilibrium**.

## Corollary

Consider the situation where there is a disease-susceptibility site and other sites in differing levels of linkage disequilibrium with the disease-susceptibility site. From large-scale genotyping or sequencing studies, we often know the matrix of pairwise *r*^2^ values, and allele frequencies at each site in the general population, broadly defined. An interesting question arises: If one has genotyped a marker site in a case/control sample set 2 and calculated 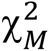 testing for disease association, can we infer the expected effect size at a non-interrogated causal site? Using **Eqn 13**, and substituting allele frequencies at the causal site,

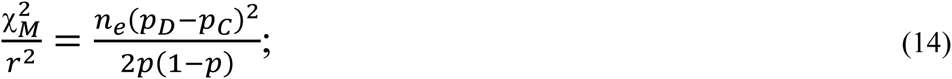

Where 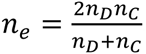 the effective total number of independent diploid samples. For an allelic odds ratio at the causal site, *R*, the allele frequency in the cases can be written as

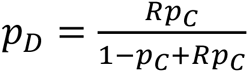

Therefore,

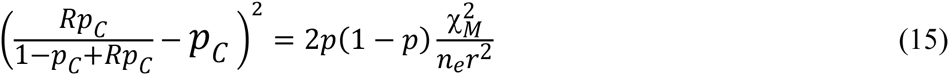

To simplify the derivation, we will assume that the disease studied is not very common such that the allele frequency in controls is well-approximated by the allele frequency in the general population, *p*_C_ ≅ *p*. This is also true if samples drawn from the general population are serving as the controls. Hence,

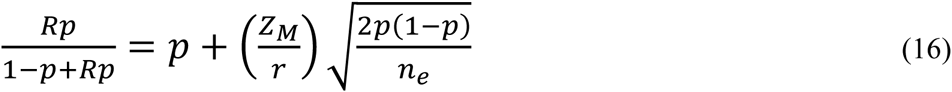

Solving for *R*,

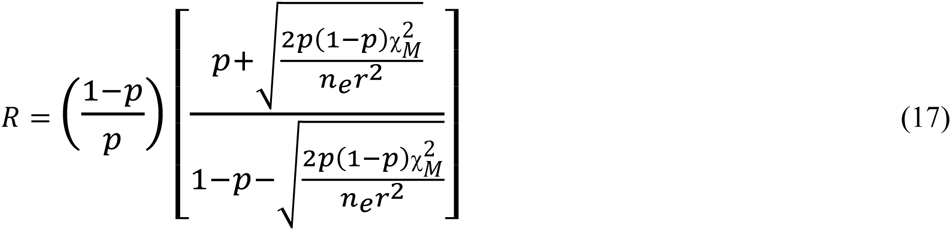

To illustrate the use and implications of **Eqn 17**, suppose that we have genotyped a site in 500 diploid cases and 500 diploid controls and calculated the test statistic χ^2^ = 20, corresponding to p-value = 7.74E-06. Further assume that this region has previously been subjected to next-generation sequencing in individuals derived from the same source population as the cases and controls which has yielded the discovery of numerous additional variants closely linked to the genotyped site, allele frequencies at those variants, and an array of pairwise linkage disequilibrium values across the region of interest. Under that scenario, one would typically have access to good estimates of the general population allele frequencies and *r*^2^ values at sites neighboring the genotyped site that produced the original finding. Suppose that one of these adjacent sites has a general population allele frequency *p* = 0.03 and a linkage disequilibrium value with the genotyped site of *r*^2^ = 0.2. Under the two-site model, we would therefore estimate the odds ratio at the putative, non-genotyped, causal site to be 5.17. Put another way, the putative causal site, having the general population allele frequency and linkage disequilibrium values above, would have to have an odds ratio of 5.17 in order to generate a Chi-Squared statistic at the genotyped site of 20 given 500 cases and 500 controls. Indirect inference of the properties of non-interrogated causal sites can be helpful in subsequent experimental efforts to identify disease-predisposing sites in a fine-mapped region. **Figure 3** displays the relationship between the inferred odds ratio at the causal site from disease association data at the marker site as a function of linkage disequilibrium between the two sites. Graphs for various *p*-values at marker site are shown.

**Figure 3.**
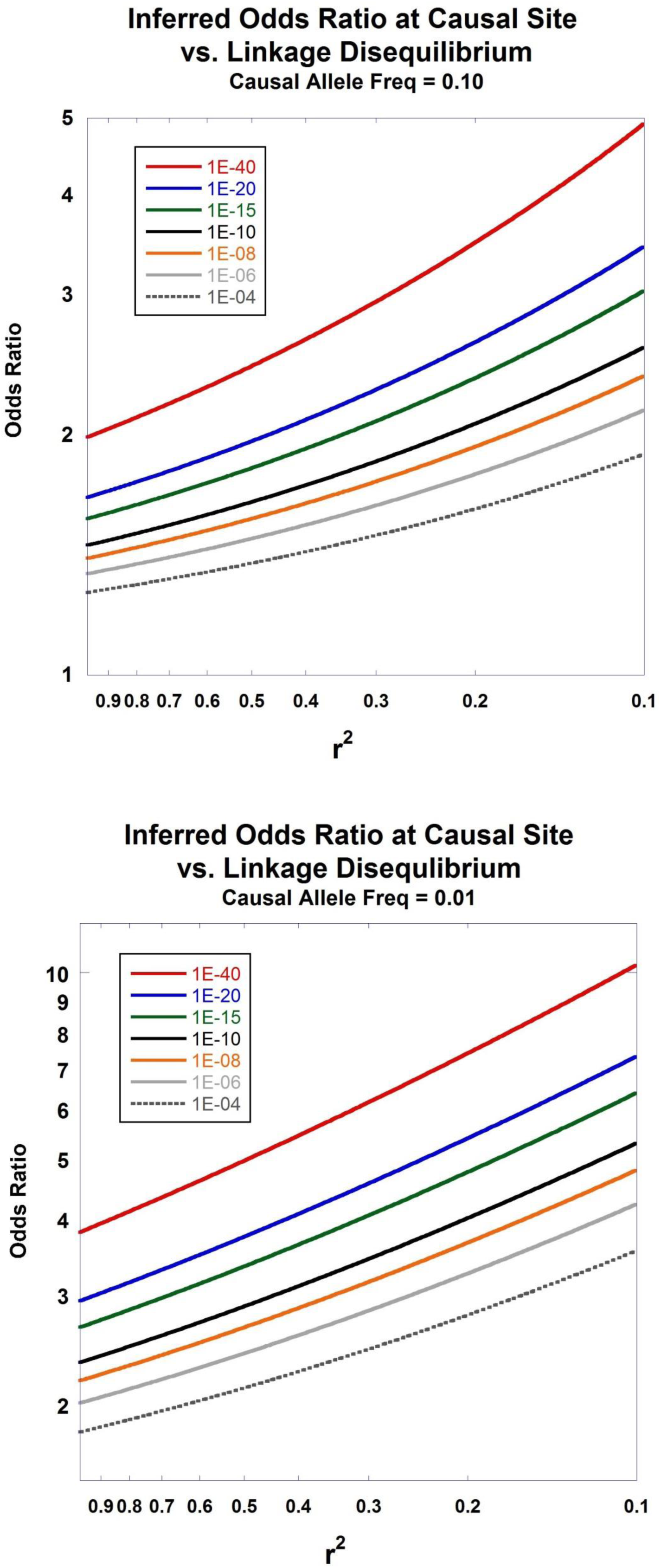
Inferred Odds Ratio. The relationship between the inferred odds ratio at a causal site and the level of linkage disequilibrium with an interrogated marker is presented in **Fig. 3a** and **Fig. 3b**. **Eqn 17** is used for the calculations. The seven curves show the patterns of expected odds ratios for disease association at the causal site under different observed p-values calculated at the marker site. Sample size was set at n_e_=5000. **Fig. 3a** shows results assuming that the disease-predisposing allele at the causal site has frequency of 0.10 in the general population, whereas **Fig. 3b** sets that frequency at 0.01.

The results detailed in **Eqns 1-17** do not treat any of the parameters, such as haplotype frequencies, as random variables. Clearly, haplotype counts in cases and controls should be treated with sampling processes from a larger population. To address this issue, we have constructed a Monte Carlo simulation program to generate haplotypes under a probabilistic model. Under this program we are able to explore the variation around **Eqn 13** and to observe effects that may be produced by different sets of parameters.

## Monte Carlo Simulations

In an effort to understand the variation in the patterns of disease association decay as a function of linkage disequilibrium with a causative site, we constructed a Monte Carlo simulation using a generalized disease model (penetrances for each of the three genotypes at the causal site are parameterized) and treating the haplotype counts in cases and controls as random variables. Recombination was introduced between a causal site and a closely linked marker as a realistic method of generating different sets of 2-site haplotypes for the general population.^39^ For a rate of recombination, *c*, and generation time *t*, we used the following set of recursions (Haldane model of recombination):

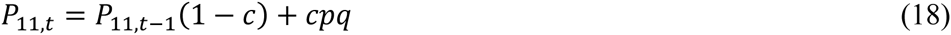

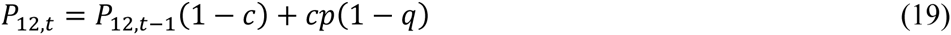

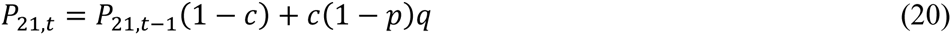

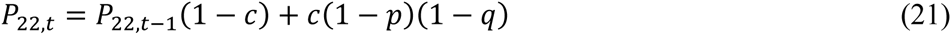

The corresponding general population allele frequency at the causal site is

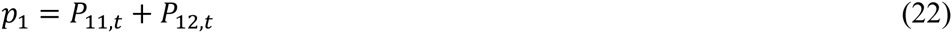

Similarly, the general population allele frequency at the linked marker is

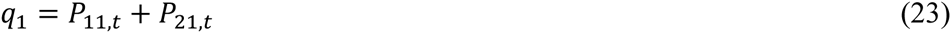

In the absence of sampling (i.e., for an infinite population size), these will be invariant under the model considered. Assuming Hardy-Weinberg equilibrium in the general population at both sites, the proportion of individuals affected by the disease attributable to this locus, is calculated through the law of total probability,

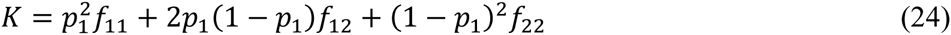

To calculate the expected haplotype frequencies in cases, we used the above general population frequencies modified through the use of Bayes theorem.^35^ Hence, the expected frequency of the *A*_1_*B*_1_ haplotype in cases is

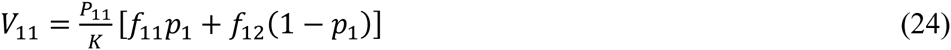

In an analogous manner, the remaining haplotype frequencies in cases, where the subscript indicates the haplotype, are

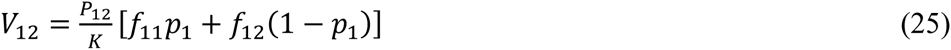

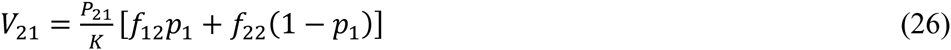

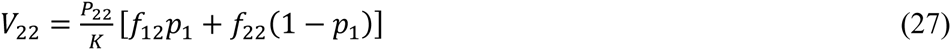

The haplotype frequencies in controls are

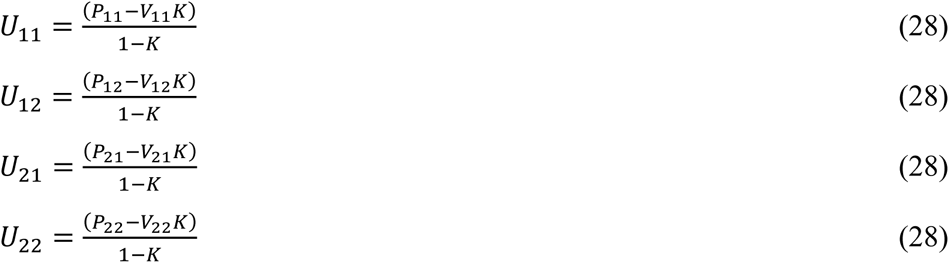

Sampling of the haplotypes from the expected frequencies is accomplished through two independent multinomial variates (one for the cases and one for the controls), such that the joint densities are given by

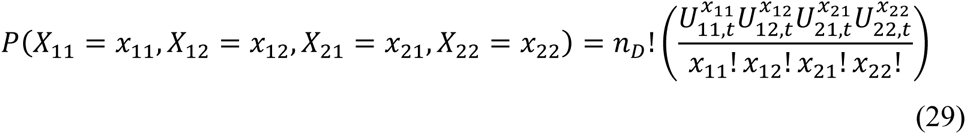

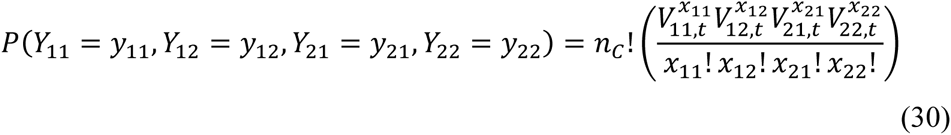

Hence, the sample frequency of the causal allele in cases and controls, respectively, are

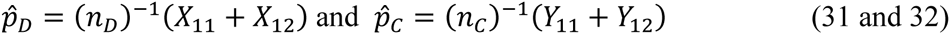

**(Plot of mean values)**

**(Plot of confidence intervals)**

## Conclusion and Discussion

One of the most fundamental patterns in disease genetics is the nature of the decay of disease association with declining linkage disequilibrium from a causal site. Motivated by the Lai et al and Pritchard-Przeworski derivations for the approximate increase in sample size to attain the equivalent statistical power at a marker site in linkage disequilibrium with a causal site, we first showed how this result could be used to produce an approximation showing a linear relationship in the Chi-Squared association statistics testing disease association at a marker and a causal site and that the ratio of the two was approximately *r*^2^ (**Eqn 2**). Next, using a general two-site model with penetrances, we showed that this is indeed an exact result and invariant to the mode of inheritance model (**Eqn 13**).

Future work focusing on imputing additional properties of a non-interrogated causal variant, or multiple causal variants, within a disease-associated region using the linkage disequilibrium patterns and disease association statistics would provide valuable insights into design and interpretation of fine mapping studies.

## Acknowledgments

We are greatly appreciative of the support from Dr. Murray Brilliant, Fritz Wenzel, Steve Ziemba, Terrie Kitchner, Cathy Marx and Marlene Stueland. This study was funded through the generous donations to the Marshfield Clinic Research Foundation and NIMH RO1 MH097464.

